# Integrated Genome Browser: visual analytics platform for genomics

**DOI:** 10.1101/026351

**Authors:** Nowlan H. Freese, David C. Norris, Ann E. Loraine

## Abstract

**Motivation:** Genome browsers that support fast navigation through vast data sets and provide interactive visual analytics functions can help scientists achieve deeper insight into biological systems. Toward this end, we developed Integrated Genome Browser (IGB), a highly configurable, interactive and fast open source desktop genome browser.

**Results:** Here we describe multiple updates to IGB, including all-new capability to display and interact with data from high-throughput sequencing experiments. To demonstrate, we describe example visualizations and analyses of data sets from RNA-Seq, ChIP-Seq, and bisulfite sequencing experiments. Understanding results from genome-scale experiments requires viewing the data in the context of reference genome annotations and other related data sets. To facilitate this, we enhanced IGB’s ability to consume data from diverse sources, including Galaxy, Distributed Annotation, and IGB-specific Quickload servers. To support future visualization needs as new genome-scale assays enter wide use, we transformed the IGB codebase into a modular, extensible platform for developers to create and deploy all-new visualizations of genomic data.

**Availability:** IGB is open source and is freely available from http://bioviz.org/igb.

## 1 Introduction

Genome browsers are visualization software tools that display genomic data in interactive, graphical formats. Since the 1990s, genome browsers have played an essential role in genomics, first as tools for building, inspecting, and annotating assemblies and later as tools for distributing data to the public (Durbin and Thierry-Mieg, 1991; Harris, 1997; Kent, et al., 2002). Later, the rise of genome-scale assays created the need for a new generation of genome browsers that could display user’s experimental data alongside reference sequence data and annotations.

Integrated Genome Browser (IGB), first developed in 2001 at Affymetrix, was among the first of this new breed of tools. IGB was first written to support Affymetrix scientists and collaborators who were using whole genome tiling arrays to probe gene expression and transcription factor binding sites as part of the ENCODE project (Kapranov, et al., 2003). As such, IGB was designed from the start to handle and display what we would now call “big data” in bioinformatics - millions of probe intensity values per sample. IGB was one of the first genome browsers to support visual analytics, in which interactive visual interfaces augment our natural ability to notice patterns in data. The “thresholding” feature described below is an example.

Because early IGB development was publicly funded, Affymetrix released IGB and its companion graphics library, the Genoviz Software Development Kit (Helt, et al., 2009), as open source software in 2004. Our first article introducing IGB appeared in 2009 (Nicol, et al., 2009) and focused on visualization of tiling array data. Here, we describe new visual analytics and data integration features developed for high-throughput sequencing data. We also introduce a new plug-in application programmers interface (API) that makes adding new functionality easier for developers, transforming IGB into an extensible visual analytics platform for genomics.

## 2 Results

IGB is implemented as a stand-alone, rich client desktop program using the Java programming language. To run IGB, users download and run platform-specific installers; these also support automatic updates. Mac and Windows installers also include a copy of the Java virtual machine, which is installed in an IGB-specific location, making it unnecessary for users to install (and maintain) Java separately.

IGB’s implementation as a local application rather than as a Web app means that IGB can access the full processing power of the user’s local computer. IGB is always present on the user’s desktop, regardless of internet connection status, but IGB can also use the Web to consume data, as described below. Because IGB runs locally, users view their own datasets without uploading them to a server, which can be important when working with confidential data.

### 3.1 Viewing genomes and annotations

On startup, IGB displays a “home” screen featuring a carousel of images linking to the latest versions of model organism, crop plant, and reference human genome assemblies. More species and genome versions are available via menus in the Current Genome tabbed panel, including more than seventy animal, plant, and microbial genomes. Users can also load and visualize their own genome assemblies, called “custom” genomes in IGB, provided they have a reference sequence file in FASTA or 2bit format.

Once a user selects a species and a genome version, IGB automatically loads the reference gene model annotations for that genome, if available. For any given genome version, additional annotations or high-throughput data sets may also be available via the Data Access Panel interface. These data sets can come from different sites, and to highlight this, IGB can display a favicon.ico graphic distinguishing these different data sources. This is why IGB is named “integrated” - it integrates data from different sources into the same view. To illustrate, Figure 1 shows an example view of integrated data sets from the human genome. Reference gene model annotations and other data sets from IGB Quickload are shown, together with tracks loaded from a DAS1 server hosted by the UCSC Genome Bioinformatics group.

**Fig. 1.**
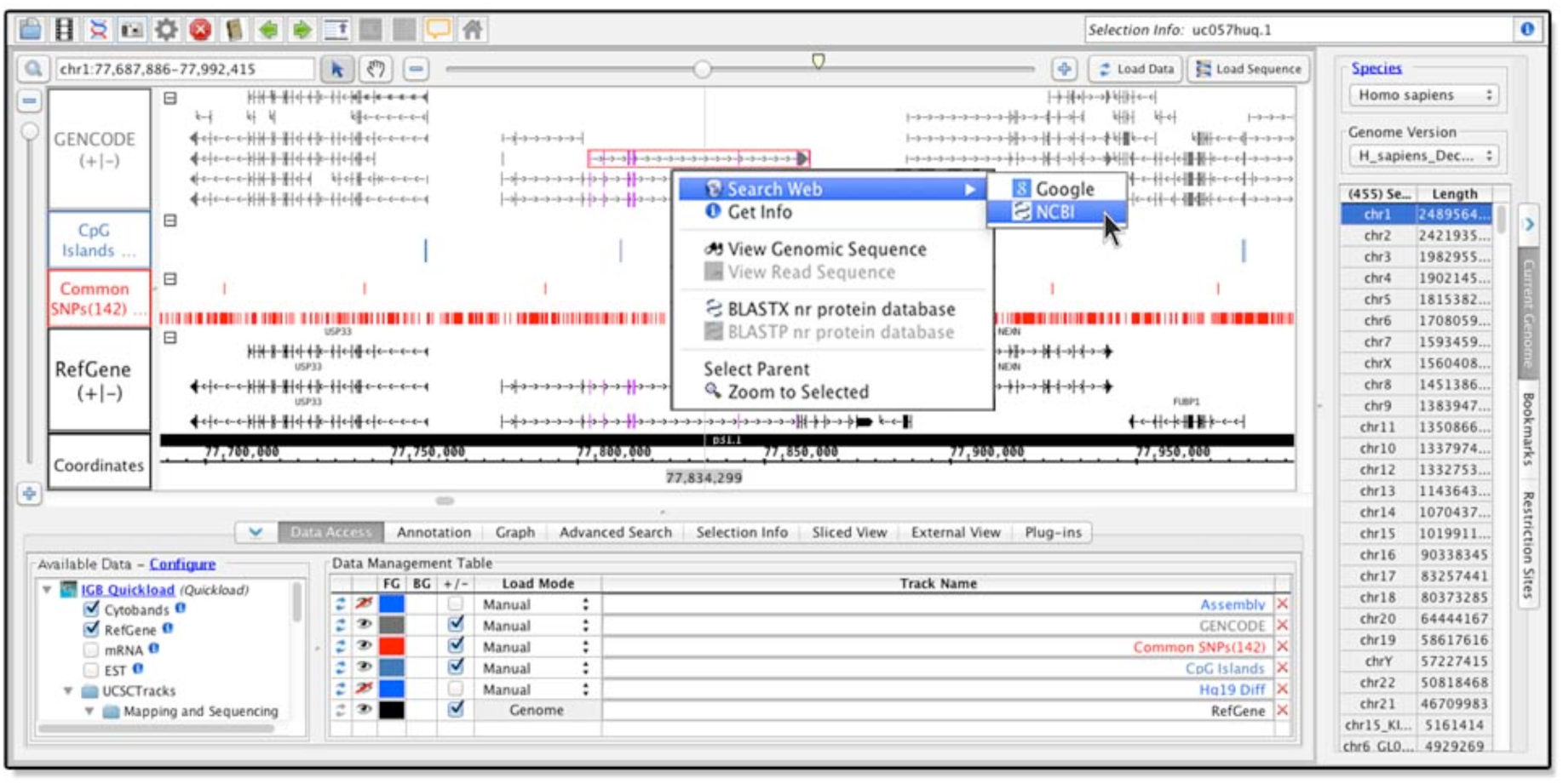
Viewing multiple human genome data sets in IGB. IGB screen showing human genome build 38, released in December 2013. Gene annotations load by defult, with additional data available in the section labeled Available Data (lower left). Each dataset occupies a separate track within the main window, and can be colored according to user preference. Right-clicking items in the main window activates a context menu with options to search Google, run BLAST, or view the underlying genomic sequence.

### 3.2 Navigating and interacting

Genomic data sets span many scales, and fast navigation through these different scales is a key feature for a genome browser. Users need to be able to quickly travel between base-level views depicting sequence details like splice sites, gene-level views depicting the exonintron organization of genes, and chromosome-level views showing larger-scale structures, like centromeres and chromosome bands. Tools that support fast navigation through the data can accelerate the discovery process.

For this reason, IGB implements a visualization technique called one-dimensional, animated semantic zooming, in which objects change their appearance in an animated fashion around a central line, called the “zoom focus” (Loraine and Helt, 2002). Animation helps users stay oriented during zooming and can create the impression of flying through one’s data (Bederson and Boltman, 1999; Cockburn, et al., 2009).

To set the zoom focus, users click a location in the display. A vertical, semi-opaque line called the “zoom stripe” indicates the zoom focus position and also serves as a pointer and guideline tool when viewing sequence or exon boundaries. When users zoom, the display appears to contract or expand around this central zoom stripe, which remains in place, thus creating a feeling of stability and control even while the virtual genomic landscape is rapidly changing.

IGB also supports “jump zooming,” in which a request to zoom triggers instantaneous teleportation to a new location. To jump-zoom, users can double-click an item, click-drag a region in the coordinates track, or search using the Advanced Search tab or the Quick Search box at the top left of the display.

Moving without changing the scale (panning) is also important for fast navigation through data. In IGB, clicking arrows in the toolbar moves the display from left to right, and scrollbars offer ways to move more rapidly. Users can click-drag the selection (arrow) cursor into the left or right border of the main display window to activate continuous pan. For finer-scale control, a move tool cursor enables click-dragging the display in any direction.

IGB helps users ask and answer questions about their data by supporting multiple ways for users to interact with what they see. Selecting data display elements (Glyphs) triggers display of meta-data about the selected item and mouse-over causes a tooltip to appear. Right-clicking items within tracks activates a context menu with options to search Google, run a BLAST search at NCBI, or open a sequence viewer (Fig. 1). Right-clicking track labels activates context menus showing a rich suite of visual analytics tools and functions, some of which we discuss in more detail below.

### 3.3 Loading data from files or URLs

IGB can consume data from local files or URLs and can read more than 30 different file formats popular in genomics. When users open a data set, a new track is added to the main display. Users then choose how much data to add to the new track by operating data loading controls. Larger data sets, such as RNA-Seq data, should be loaded on a region-by-region basis, while others that are small enough to fit into memory can be loaded in their entirety, e.g., a BED file containing ChIP-Seq peaks. To load data into a region, users zoom and pan to the region of interest and click a button labeled “Load Data.” To load all data in an opened data set, they can change the data sets loading method by selecting a “genome” load mode setting in the Data Access Panel.

This behavior differs from other genome browsers in that most other tools link navigation and data loading. In other tools, a request to navigate to a new location or change the zoom level both redraws the display and also triggers a data loading operation. Although sometimes convenient for users, this limits the types of navigation interactions a tool can support. This trade-off can be seen in the Integrative Genomics Viewer (IGV), a desktop Java application developed after IGB (Thorvaldsdottir, et al., 2013). IGV auto-loads data but restricts movement; for example, it lacks panning scrollbars and does not support fast, animated zooming. IGB, by contrast, prioritizes navigation speed and gives users total control over when data load, which can be important when loading data from distant locations over slower internet connections. IGB gives users control over when such delays might occur, thus making waiting more palatable.

### 3.4 Sharing and integrating data

IGB aims to support the scientific discovery process by making it easy for users to document and share results. Taking a cue from Web browsers, IGB for many years has supported bookmarking genomic scenes. In IGB, genomic scene bookmarks record the location, genome version, and data sets loaded into the current view. Users can also add free text notes and thus record conclusions about what they see. Selecting a bookmark causes IGB to zoom and pan to the book-marked location and load all associated data sets, thus enabling users to quickly return to a region of interest. Users can sort, edit, import and export bookmarks using the Bookmarks tab. Exporting bookmarks creates an HTML file which users can re-import into IGB or open in a Web browser. If IGB is running, clicking an IGB bookmark in a Web browser causes IGB to zoom to the bookmarked location and load associated data.

IGB loads bookmarks through a ReST-style endpoint implemented within IGB itself, using a port on the user’s computer. This endpoint allows bookmarks to be loaded from HTML hyperlinks embedded in web pages, spreadsheets, or any other document type that supports hyperlinks. IGB was the first desktop genome browser to implement this technique; for many years, Affymetrix used it to display probe set alignments on their NetAffx Web site (Liu, et al., 2003).

More recently, we used IGB’s ReST-style bookmarking system to implement a Javascript bridge between IGB and the Galaxy bioinformatics workflow system (Goecks, et al., 2010). When users generate IGB-compatible data files within Galaxy, they can now click a ‘display in IGB’ hyperlink. Clicking this link opens a BioViz.org Web page containing a javascript program that forwards the hyperlink to IGB, causing IGB to retrieve that data directly from Galaxy. If IGB is not running, the javascript instead invites the user to launch IGB. Once they do, the data set loads.

IGB supports multiple formats and protocols for sharing data set collections and integrating across data sources. The most lightweight and easy to use of these is the IGB specific “Quickload” format, which consists of a simple directory structure containing plain text, metadata files. The metadata files can reference data sets stored in the same Quickload directory or in other locations, including Web, ftp sites, or cloud storage resources such as Dropbox and iPlant (Goff, et al., 2011). This flexibility makes it possible for a Quickload site to aggregate data sets from multiple locations. The metadata for a Quick-load site also includes styling and data access directives controlling the way IGB loads each file, and how the data will look when loaded.

Sharing a Quickload site is straightforward. Users simply copy the contents to a publicly accessible location and then publicize the URL, which IGB users then add as a new Quickload site to their copy of IGB. Typically, users copy their Quickload sites to the content directories of Web sites, and IGB supports secure access via the HTTP basic authentication protocol, making it easy to keep data sets private if desired. If users do not require password-protection for their data, they can also use Public Dropbox folders to share Quickload sites.

### 3.5 Visualizing RNA-Seq data

As ultra high-throughput sequencing of cDNA (RNA-Seq) has supplanted microarrays for surveying gene expression, we added new features to IGB to support visualization of RNA-Seq data. RNA-Seq analysis data processing workflows typically produce large read alignment (BAM) files, which IGB can open and display. IGB also implements many new visual analytics functions that operate on BAM file tracks and highlight biologically meaningful patterns in the data.

Right-clicking a BAM track label opens a context menu listing options to create coverage graphs, called “depth” graphs in IGB. At present, IGB supports two depth graph types: “depth graph start” that counts a read’s first mapped base and “depth graph all” that counts the number of reads overlapping a position. The latter is useful for view ing overall expression at a locus, and the former is useful for investigating sequencing bias. Both graph types are implemented using IGB’s track operations API, which developers can use to add all-new graph generation algorithms as plug-ins, described in section 3.8.

By comparing depth graphs for multiple samples, users can identify differences in transcript levels. Using the Graph tab, users can place multiple graphs on the same scale, and if sequencing depth is similar, peaks that are the same height and shape reflect similar expression. Figure 2a shows “depth graph all” graphs made from RNA-Seq alignment (BAM) tracks; reads were from sequencing lung cancer samples bearing wild-type or mutant copies of the *KRAS* oncogene (Kalari, et al., 2012). One peak is much taller in the mutant sample, indicating higher expression. The gene is *AQP3*, encoding a water/glycerol-transporting protein (Hara-Chikuma and Verkman, 2005).

**Fig. 2.**
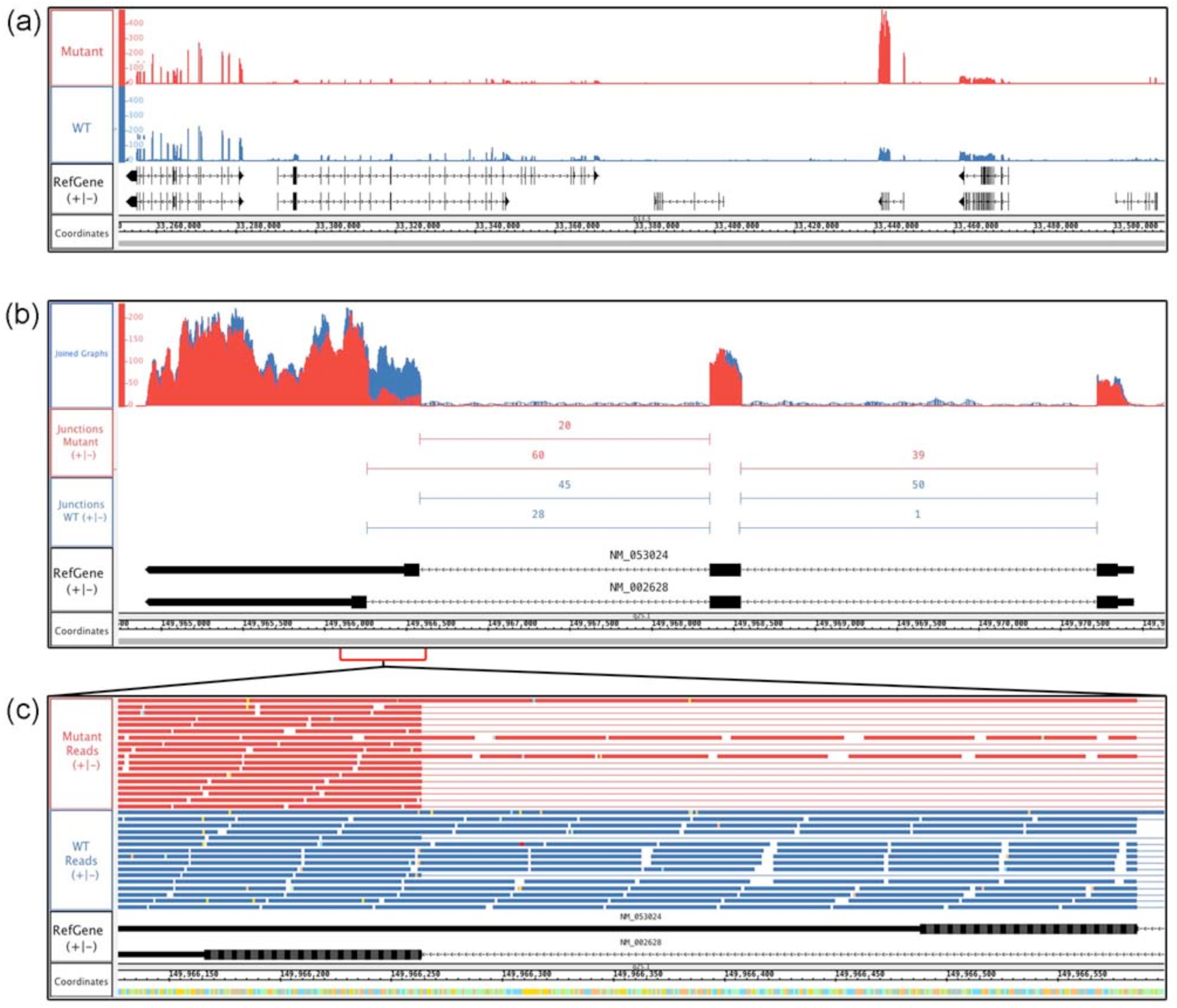
RNA-Seq data from human lung adenocarcinomas bearing mutant or wild-type (WT) alleles of the KRAS oncogene. (**a**) Coverage depth graphs show transcript abundance across a 250 kb region. Mutant samples contain a peak indicating higher expression in the mutant sample. **(b)** Overlaid depth graphs showing a discontinuity in coverage indicating differential splicing in *PFN2*. Quantification of split reads by FindJunctions further supports differential splicing. **(c)** Zoomed in view of (b), showing aligned reads.

Visual analysis of RNA-seq data can also highlight alternative splicing differences between samples. Figure 3b shows *PFN2*, which produces two isoforms due to alternative splicing. Here, the depth graphs from Figure 2a were merged into a single track. Comparing the peak discontinuities to the gene models shows that the mutant sample favors the shorter isoform. To further aid splicing analysis, we developed FindJunctions, a visual analytics tool that identifies split read alignments in an RNA-Seq track, uses them to identify potential exon-exon junctions, and then creates an all-new track containing junction features annotated with the number of alignments that supported them. Figure 2b shows an example where FindJunctions split read quantification of split reads reinforces the finding that *PFN2* is differentially spliced between samples. Viewing the read alignments is also informative. Figure 2c shows a zoomed-in view of the alternatively spliced region from Fig. 2b. Read alignments show that the shorter isoform predominates in the sample bearing mutant *KRAS*.

**Fig. 3.**
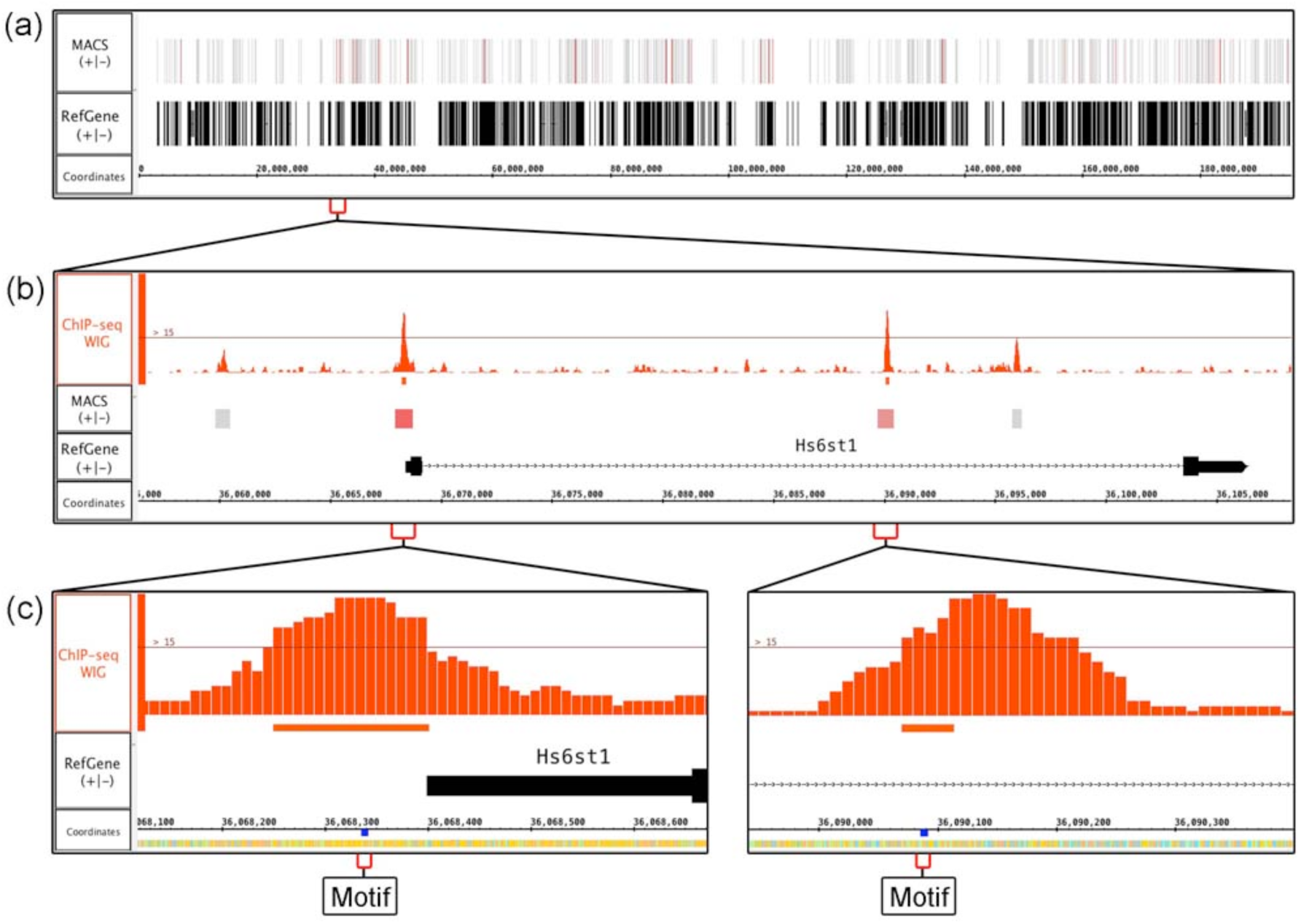
Visualizing ChIP-Seq data. (**a**) MACS BED file with peak regions from mouse ChIP-Seq data investigating binding sites for transcription factor SOX9. Peak regions in the track labeled “MACS” are colored by score, and higher-scoring regions appear more red. **(b)** Zoomed in view of (a). MACS identified four significant peaks, of which two exceeded a user-defined coverage threshold, visible as a thin horizontal line in the ChIP-Seq WIG track (top). **(c)** Zoomed in view of (b). Searching for the motif AGCCGYG identified sites under the most significant peaks in (b).

Users can interact with reads and other data displayed in the viewer using selection operations. Clicking a single item selects it, and click-dragging over multiple items selects all of them. Pressing keyboard modifiers while clicking an item adds (SHIFT) or removes (CNTRL) it from the pool of selected items. IGB reports the identity or number of selected items in the upper right corner. In this way, users can identify and count items that match a biologically interesting pattern, e.g., spliced reads that support a junction. As an additional visual cue, when a read or annotation is selected, all items with matching boundaries on either the 5’ or 3’ end are highlighted. This technique, called “edge matching,” aids in visualization of read boundaries and identifying alternative donor/acceptor sites.

### 3.6 Visualizing ChIP-Seq data

Knowing where a transcription factor binds DNA in relation to nearby genes is tantamount to understanding its function, as genes whose promoters are bound by a given transcription factor are likely to be regulated by it. Identifying binding sites of DNA-binding proteins is now routinely done using whole-genome ChIP-Seq, in which DNA cross-linked to protein is immunoprecipitated using antibodies against the protein of interest and then sequenced. Subsequent data analysis typically involves mapping reads onto a reference genome, identifying regions with large numbers of immunoprecipated reads, and then performing statistical analysis to assess significance of enrichment. Each step requires users to choose analysis parameters whose effects and importance may be hard to predict. To validate these choices, it is important to view the “raw” alignments and statistical analysis results in a genome browser. As an example, we describe using IGB to view results from a ChIP-Seq analysis done using MACS, a widely used tool (Zhang, et al., 2008).

MACS produces a BED format file containing peak locations and significance scores indicating which peaks likely contain a binding site. Opening and loading this BED file in IGB creates a new track with single-span annotations representing the extent of each peak. At first, they look identical, varying only by length, and it is difficult to distinguish them. To highlight the highly scoring peak regions, IGB offers a powerful “Color by” visual analytics feature that can assign colors from a heatmap using quantitative variables associated with features. For this, users right-click a track, select “Color by” from the context menu, and then operate a heatmap editor adopted from the Cytoscape codebase to color-code by score (Shannon, et al., 2003). Doing this makes it easy to identify the highest scoring region most likely to contain a binding site (Fig. 3a).

ChIP-Seq analysis tools also typically produce WIG-format depth-graph files that report where immunoprecipitated sequences have “piled up,” forming peaks. Loading this WIG file into IGB creates a new graph track that shows how these peaks coincide with regions from the BED file. In IGB, users can move tracks to new locations in the display. As shown in Figure 4b, placing the WIG track above the color-coded BED track makes it easy to observe how taller peaks typically have higher significance values.

**Fig. 4.**
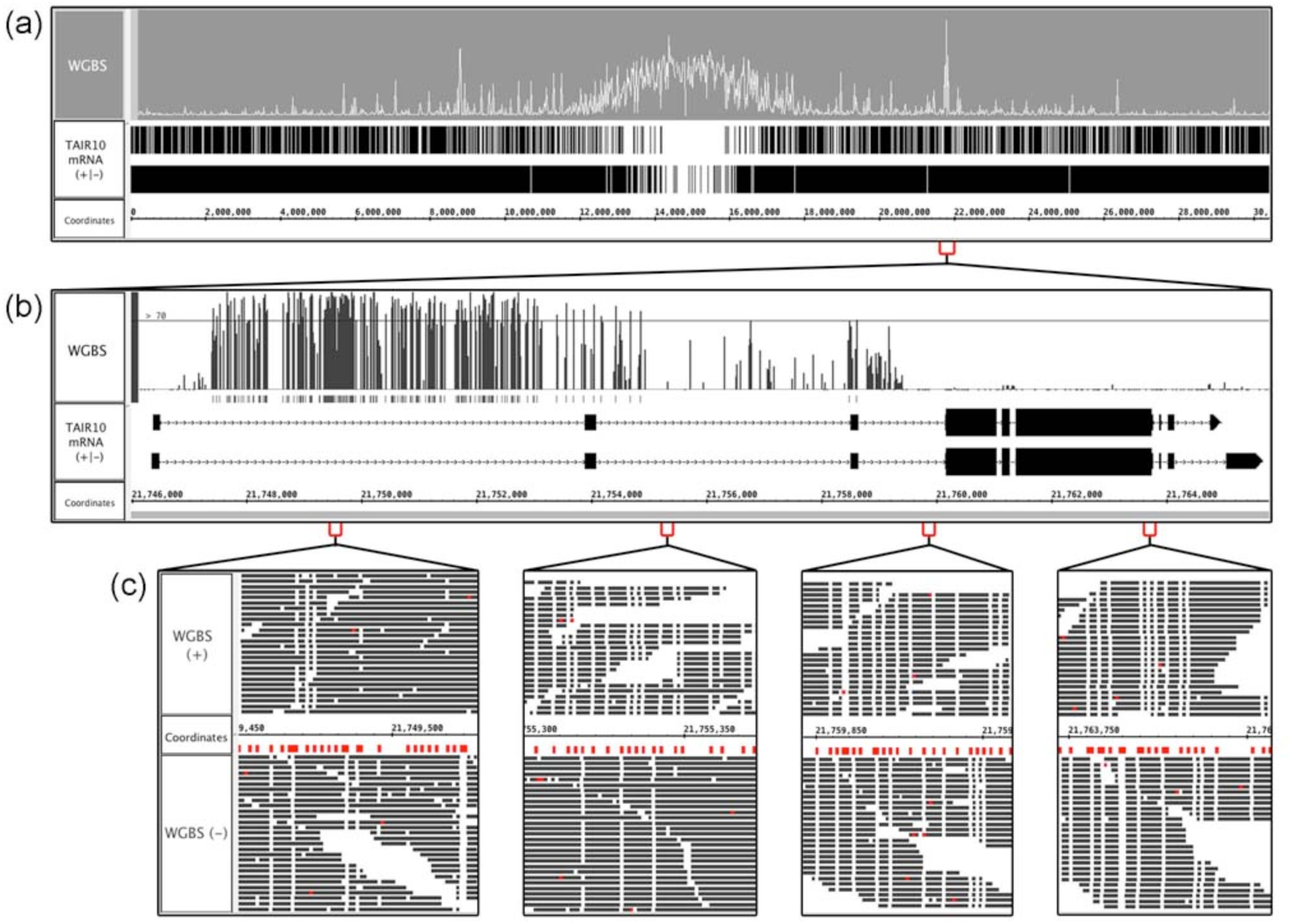
Visualizing bisulfite sequencing data. (**a**) Bismark bedGraph file from an *Arabidopsis* bisulfite sequence experiment. Peaks indicate regions containing many methylated cytosine residues. **(b)** Zoomed in view of (a). A user-defined threshold shows that most cytosines in the first intron are methylated. **(c)** Zoomed in view of (b), showing the positive and negative strand aligned reads. Thymines are colored white, and cytosines red. Unmethylated cytosines that were converted to thymines appear as white columns occupying the same base pair position as a mark below the sequence axis, which indicate cytosines in the reference sequence. The highly-methylated region on the left contains many marks and few white columns.

IGB’s graph thresholding function, originally developed for tiling arrays, is useful for exploring the relationship between coverage depth and peak score. Available from the Graph tab, thresholding identifies regions in a graph track where consecutive y-values exceed a user-defined value. Users can change this threshold value dynamically using a slider and observe in real-time how this affects the number and extent of identified regions. Users can promote regions to new tracks and save them in BED format. As an example, Figure 4b shows four MACS-detected peaks near the *Hs6st1* gene, a regulatory target for the SOX9 transcription factor in mouse (Kadaja, et al., 2014). Two peaks exceed the threshold, providing a visual cue that these locations may be most important for regulation.

Sometimes the recognition sequence for transcription factor being studied is known or can be deduced from the data. IGB offers a way for users to visualize instances of binding site motifs. Using the Advanced Search tab, users can search the genomic region in view using regular expressions. For instance, to search for the motif AGCCGYG (where Y can be C or T) a user would enter “AGCCG[CT]G”. In this example, the search found several instances of this motif, with one located in each of the two user-defined significant peaks near the *Hs6st1* gene (Fig. 3c).

### 3.7 Visualizing bisulfite sequencing data

Whole genome bisulfite sequencing (WGBS) refers to bisulfite conversion of unmethylated cytosines to thymines followed by sequencing. This technique can identify methylated sites throughout a genome and reveal potential epigenetic regulation of gene expression. Analyzing bisulfite data involves mapping the reads onto the genome using tools that can accommodate reads where many but not all cytosine residues have been converted. Several such tools are available; here we describe using IGB to visualize output of Bismark (Krueger and Andrews, 2011).

Bismark produces a BAM file containing read alignments and a depth-graph file (in bedGraph format) reporting percent methylation calculated from sliding windows along the genome. Figure 4a shows a Bismark bedGraph file from an experiment investigating methylation in the model plant *Arabidopsis thaliana* (Yelagandula, et al., 2014). All of chromosome one is shown, and peaks indicate regions of high methylation. This whole chromosome view makes it easy to observe that the centromeric region is highly methylated. From here, users can zoom in to examine methylation at a region, gene, or base pair level.

Figure 4b shows a zoomed-in view of one of the tallest peaks visible in Figure 4a. Similar to the ChIP-Seq data analysis described in the previous section, users can apply the thresholding feature to identify regions of high methylation within a gene. In this example, the threshold for percent methylation is set to 70% or greater, highlighting how the first intron, but not the second and third introns, is highly enriched with methylated cytosines.

Closer examination of aligned reads provides further support of methylation. In IGB, nucleotide residues are color-coded, and users can change these colors. In addition, read alignment tracks can be configured so that only mismatched bases are color-coded. Figure 4c shows a view of bisulfite sequencing data in which cytosines are red and thymines are white. In this view, white columns in the read track (labeled WGBS) represent unmethylated cytosines, and marks in the coordinates track indicate cytosine residues in the reference sequence that. Regions with many cytosines and few white columns are highly methylated.

### 3.8 Extending IGB using plug-ins

Developers have created dozens of genome browser tools, each one aiming to meet a need not met by the tools that preceded it. And yet, each new tool has faced similar problems, such as how to consume data from files, how to lay out genomic features and graphs into tracks, and how to support zooming through vast differences in scale. As described above, IGB has solved many of these problems and offers a flexible and fast environment for users to explore the genomic landscape. In order for developers to quickly create new visualizations we transformed IGB into a modular, extensible platform for developers to create and deploy all-new visualizations of genomic data.

The IGB software architecture now resembles other popular open source Java-based projects that support adding new functionality via plug-ins, including Eclipse, the Netbeans Rich Client Platform, and Cytoscape (Shannon, et al., 2003). This similarity comes from our common use of OSGi, a services based architectural framework and community standard for building modular software. By adopting this framework, we increased IGB extensibility, simplified adding new features, and created a plug-in API that empowers community developers to contribute new functionality without needing deep understanding of IGB internal systems. The plug-in API is new, but community developers are already using it to create novel visualizations (Céol and Müller, 2015). Documentation describing how to create plug-ins for IGB is available from the IGB developer’s guide.

## 3 Future directions

IGB offers users powerful utilities for viewing, analyzing, and interacting with data within an environment that feels fast, flexible, and highly interactive. In the future, we plan to make the IGB’s Quickload data sharing system easier to use by providing tools for building Quickload sites from within IGB, along with a Quickload registry for users to publicize their sites. In addition, we will continue developing and improving the IGB plug-in APIs, providing documentation and example plug-ins to demonstrate how developers can use IGB as a platform to support genomics research.

## Acknowledgements

We thank the many developers, testers, and designers who contributed to IGB, some of whom include Gregg Helt, Michael Lawrence, Lance Frohman, John Nicol, Hiral Vora, Alyssa Gulledge, Fuquan Wang, David Nix, Ido Tamir, Vikram Bishnoi, Anuj Puram, Richard Linchangco, Mason Myer, Katherine Kubiak, Zhong Ren, Tarun Kanaparthi, Kyle Suttlemyre, and Tarun Mall.

## References

Bederson, B.B. and Boltman, A. (1999) Does animation help users build mental maps of spatial information? In, Information Visualization. (Info Vis ’99) Proceedings. IEEE Symposium. p. 28–35.

Céol, A. and Müller, H. (2015) The MI Bundle: Enabling Network and Structural Biology in genome visualization tools. Bioinformatics.

Cockburn, A., Karlson, A. and Bederson, B.B. (2009) A review of overview+detail, zooming, and focus+context interfaces. ACM Comput. Surv. 3;41(1):1–31.

Durbin, R. and Thierry-Mieg, J. (1991) A C. elegans Database. In. Documentation, code and data available from anonymous FTP servers at lirmm.lirmm.fr, cele.mrc-lmb.cam.ac.uk and ncbi.nlm.nih.gov.

Goecks, J., Nekrutenko, A. and Taylor, J. (2010) Galaxy: a comprehensive approach for supporting accessible, reproducible, and transparent computational research in the life sciences. Genome Biol 3;11(8):R86.

Goff, S.A., et al. (2011) The iPlant Collaborative: Cyberinfrastructure for Plant Biology. Front Plant Sci 3;2:34.

Hara-Chikuma, M. and Verkman, A.S. (2005) Aquaporin-3 functions as a glycerol transporter in mammalian skin. Biology of the cell / under the auspices of the European Cell Biology Organization 3;97(7):479–486.

Harris, N.L. (1997) Genotator: a workbench for sequence annotation. Genome Res 3;7(7):754–762.

Helt, G.A., et al. (2009) Genoviz Software Development Kit: Java tool kit for building genomics visualization applications. BMC bioinformatics 3;10:266.

Kadaja, M., et al. (2014) SOX9: a stem cell transcriptional regulator of secreted niche signaling factors. Genes Dev 3;28(4):328–341.

Kalari, K.R., et al. (2012) Deep Sequence Analysis of Non-Small Cell Lung Cancer: Integrated Analysis of Gene Expression, Alternative Splicing, and Single Nucleotide Variations in Lung Adenocarcinomas with and without Oncogenic KRAS Mutations. Frontiers in Oncology 3;2:12.

Kapranov, P., Sementchenko, V.I. and Gingeras, T.R. (2003) Beyond expression profiling: next generation uses of high density oligonucleotide arrays. Briefings in functional genomics & proteomics 3;2(1):47–56.

Kent, W.J., et al. (2002) The human genome browser at UCSC. Genome Res 3;12(6):996–1006.

Krueger, F. and Andrews, S.R. (2011) Bismark: a flexible aligner and methylation caller for Bisulfite-Seq applications. Bioinformatics 3;27(11):1571–1572.

Liu, G., et al. (2003) NetAffx: Affymetrix probesets and annotations. Nucleic Acids Res 3;31(1):82–86.

Loraine, A.E. and Helt, G.A. (2002) Visualizing the genome: techniques for presenting human genome data and annotations. BMC bioinformatics 3;3:19.

Nicol, J.W., et al. (2009) The Integrated Genome Browser: free software for distribution and exploration of genome-scale datasets. Bioinformatics 3;25(20):2730–2731.

Shannon, P., et al. (2003) Cytoscape: a software environment for integrated models of biomolecular interaction networks. Genome Res 3;13(11):2498–2504.

Thorvaldsdottir, H., Robinson, J.T. and Mesirov, J.P. (2013) Integrative Genomics Viewer (IGV): high-performance genomics data visualization and exploration. Brief Bioinform 3;14(2):178–192.

Yelagandula, R., et al. (2014) The histone variant H2A.W defines heterochromatin and promotes chromatin condensation in Arabidopsis. Cell 3;158(1):98–109.

Zhang, Y., et al. (2008) Model-based analysis of ChIP-Seq (MACS). Genome Biol 3;9(9):R137.

